# Mutations in SIX1 associated with Branchio-oto-renal Syndrome (BOR) differentially affect otic expression of putative target genes

**DOI:** 10.1101/2021.06.11.447956

**Authors:** Tanya Mehdizadeh, Himani Datta Majumdar, Sarah Ahsan, Andre Tavares, Sally A. Moody

## Abstract

Single nucleotide mutations in *SIX1* are causative in some individuals diagnosed with branchio-otic/branchio-oto-renal (BOR) syndrome. To test whether these mutations have differential effects on otic gene expression, we engineered four BOR mutations in *Xenopus six1* and targeted mutant protein expression to the neural crest and cranial placode precursor cells in wild-type embryos. Changes in the otic expression of putative Six1 targets and/or co-factors were monitored by qRT-PCR and in situ hybridization. We found that each mutant had a different combination of effects. The V17E mutant reduced *eya2, tspan13, zbtb16* and *pa2g4* otic vesicle expression at a frequency indistinguishable from wild-type Six1, but reduced *prdm1* more and *spry1* less compared to wild-type Six1. For most of these genes, the R110W, W122R and Y129C mutants were significantly less repressive compared to wild-type Six1. Their individual effects varied according to the level at which they were expressed. The R110W, W122R and Y129C mutants also often expanded *prdm1* otic expression. Since previous studies showed that all four mutants are transcriptionally deficient and differ in their ability to interact with co-factors such as Eya1, we propose that altered co-factor interactions at the mutated sites differentially interfere with their ability to drive otic gene expression.

## INTRODUCTION

Branchio-otic/Branchio-oto-renal (BOR) syndrome is an autosomal dominant developmental disorder responsible for a variable combination of developmental defects including hyoid fistulas/cysts, inner, middle and external ear malformations leading to conductive and/or sensorineural hearing loss, and in some cases kidney dysmorphology [1–3]. Point mutations that result in single amino acid substitutions have been identified in about half of BOR cases, including SIX1, a homeodomain-containing transcription factor (~4% of patients), and EYA1, a co-factor that binds to SIX1 to modify its transcriptional activity (~ 40% of patients) [3,4]. Like other members of the SIX family, SIX1 contains two highly conserved domains: a SIX-type homeodomain (HD) that binds to DNA and an N-terminal SIX domain (SD) that binds to co-factor proteins that modify its transcriptional activity [5–8]. All of the reported *SIX1* mutations in BOR are located in either the SD or the HD [9–14]. Studies in several vertebrates indicate that *Six1* is a crucial regulator of cranial placode development [15–19]. Six1 loss-of-function studies demonstrate reduced expression of placode genes and/or defects in otic development [20–33]. Conversely, *Six1* gain-of-function studies show that expansion of placode domains at the expense of the adjacent epidermal, neural crest and neural plate regions [21,29,34,35].

In BOR, there is considerable variability in the presence, type and severity of the clinically relevant abnormalities even among family members carrying the same mutation, making it difficult to correlate the type of mutation with the clinical presentation [9–12, 36,37]. Therefore, we introduced four of the human mutations into *Xenopus Six1* [14] to address whether some of the variability might be attributed to changes in gene expression in the embryonic precursors of the affected cranial tissues. The mutations include: a V17E mutation in the first α-helix of the SD that interferes with the SIX1-EYA interaction and subsequent translocation of EYA to the nucleus; an R110W mutation in the sixth α-helix of the SD that decreases the EYA/SIX interaction and reduces transcriptional activation; a W122R mutation adjacent to the N-terminus of the HD that disturbs interactions with EYA1 and is transcriptionally deficient; and a Y129C mutation in the N-terminal region of the HD that significantly reduces DNA binding and transcriptional activity [10, 12–14]. We found that expressing each Six1 mutant protein in a wild-type embryo had different effects on neural border, neural crest, pre-placodal ectoderm (PPE) and a few otic vesicle genes [14]. In addition, morphological assessment of the tadpole inner ear demonstrated that the auditory and vestibular structures formed, but the volume of the lumen, the otic capsule, otoconia, and some sensory hair cell end-organs were differentially affected [14].

Although changes in otic gene expression and morphology are altered by these BOR Six1 mutant proteins, we do not know whether these changes are the direct consequence of altering the expression of Six1 target genes. Large-scale screens in flies and vertebrates have identified hundreds of Six1 transcriptional targets in several tissues and developmental stages [38–42]. Therefore, to determine whether Six1 BOR mutations alter the expression of putative transcriptional targets, we chose five putative targets based on these previous studies (*eya2, prdm1, spry1, tspan13, zbtb16*), as well as one novel cofactor (*pa2g4*) [43]. Specifically, *eya2* is expressed in the pre-placodal region, acts as a Six1 co-factor to promote pre-placodal gene expression and suppress neural plate and neural crest cell fates [29,44,45]. *prdm1* (aka *blimp1*) encodes a transcription factor that plays a role in pharyngeal arch, neural crest and placode formation [46–49]. *spry1* encodes a negative regulator of FGF signaling and is expressed in several craniofacial tissues [50–52]. *tspan13*, a member of the *tetraspanin* gene family, encodes a cell-surface protein that mediates signal transduction events in developing systems and cancers [53]. *zbtb16* encodes a Krueppel C2H2-type zinc finger protein expressed in craniofacial and several other developing tissues [54,55]. *pa2g4* (aka *ebp1*) is expressed in craniofacial tissues and cancers [56–58], and its protein binds to Six1 to regulate the development of neural crest and PPE [43].

By expressing four different mutant protein in wild-type embryos, we found that each of them reduced *spry1* expression moderately less than wild-type Six1. For *eya2, tspan13, zbtb16* and *pa2g4*, the V17E mutant reduced expression at a frequency indistinguishable from wild-type Six1, whereas the R110W, W122R and Y129C mutants were significantly less repressive. V17E reduced *prdm1* expression more frequently than wild-type Six1, whereas the R110W, W122R and Y129C mutants reduced it less frequently than wild-type Six1 and often expanded its otic expression. Since previous studies showed that each of the four mutant proteins are transcriptionally deficient and have different abilities to interact with co-factors such as Eya1, we propose that altered co-factor interactions at the mutated sites differentially interfere with their ability to drive otic gene expression.

## MATERIALS AND METHODS

### Obtaining embryos and microinjections

Fertilized *Xenopus laevis* embryos were obtained by gonadotropin-induced natural mating of adult frogs [59]. Embryos were picked at the 2-cell stage to accurately identify the dorsal and ventral animal blastomeres [59–61]. When these embryos reached the 8-cell stage, the dorsal-animal and ventral-animal blastomeres that extensively give rise to the neural crest and cranial placodes [62] were microinjected on one side of the embryo as previously described [59], with 1nl of: 1) one of the *Six1* mRNAs (wild type, fusion protein, or mutant), mixed with β-galactosidase mRNA as a lineage tracer; or 2) a 1:1 mixture of two lissamine-tagged antisense morpholino oligonucleotides that previously were verified to effectively and specifically block the translation of endogenous Six1 (Six1-MOs) [21]. Following, embryos were cultured in a dilution series of Steinberg’s medium until fixation.

### In vitro synthesis of mRNAs and antisense RNA probes

Transcripts encoding Six1 wild-type (Six1WT) [21], Six1WT SD+HD fused to the VP16 activation domain (Six1VP16) [21], Six1WT SD+HD fused to the En2 repressive domain (Six1EnR) [21], four different Six1 mutants (V17E, R110W, W122R, Y129C) [14] and a nuclear-localized βgalactosidase mRNA (as a lineage tracer) were synthesized *in vitro* according to manufacturer’s protocols (mMessage mMachine Kit, Ambion). Plasmids encoding *eya2* (Open Biosystems), *prdm1* (Open Biosystems), *spry1* (Dharmacon), *tspan13* (Dharmacon), *zbtb16* (Source Bioscience) and *pa2g4* (Open Biosystems) were purchased and sequenced in both directions to confirm identity. Digoxigenin-labeled antisense RNA probes for *in situ* hybridization (ISH) assays were synthesized from these plasmids *in vitro* according to manufacturer’s protocols (MEGAscript Kit; Ambion).

### Histochemistry, in situ hybridization (ISH) and analyses

Embryos were cultured to otic pit (st 20-22) or otic vesicle (st 26-32) stages [63], fixed in 4% paraformaldehyde (in 0.1M MOPS, 2mM EGTA Magnesium, 1mM MgSO_4_, pH 7.4), stained for β-Gal histochemistry in those embryos injected with mRNA, and processed for ISH according to standard protocols [64]. The analysis included only those embryos in which either the β-Gal-positive (mRNA-injected) or lissamine-labeled (MO-injected) cells were located in the otic region, indicating that the injected reagent was targeted to the correct tissue. In each embryo, the intensity and size of the otic expression domain of each gene was compared between the injected, lineage-labeled side of the embryo to the uninjected, control side of the same embryo, providing paired analyses. Assays were repeated a minimum of three times on embryos derived from three different sets of outbred parents. At least two authors independently scored embryos for gene expression changes. Differences in the frequency of gene expression changes were tested for significance (p<0.05) by the chi-square test (GraphPad Prism software).

### qPCR analyses

Both cells of 2-cell embryos were injected with either *Six1WT* mRNA (150pg or 400pg/cell) or mutant mRNAs (V17E, 150pg; R110W 400pg; W122R 400pg; Y129C 400pg/cell), grown to stage 32 and ten dissected heads were collected in TRI-reagent (Zymo) and processed for RNA extraction with DNAse I treatment using the Direct-zol RNA Miniprep kit (Zymo). cDNA was synthesized using the iScript Advanced cDNA Synthesis kit (Bio-Rad). qPCR was performed using 5ng cDNA with the SsoAdvanced Universal SYBR Green Mix (Bio-Rad). Primer sequences are listed in Table 1. qPCR of three biological replicates (ten heads per replicate) was performed in duplicate. PCR and data analysis were performed using a CFX Connect thermocycler (Bio-Rad). Statistical analysis was performed with GraphPad Prism 9, with significance calculated by two-way ANOVA followed by Tukey’s multiple comparisons test.

**Table 1.**
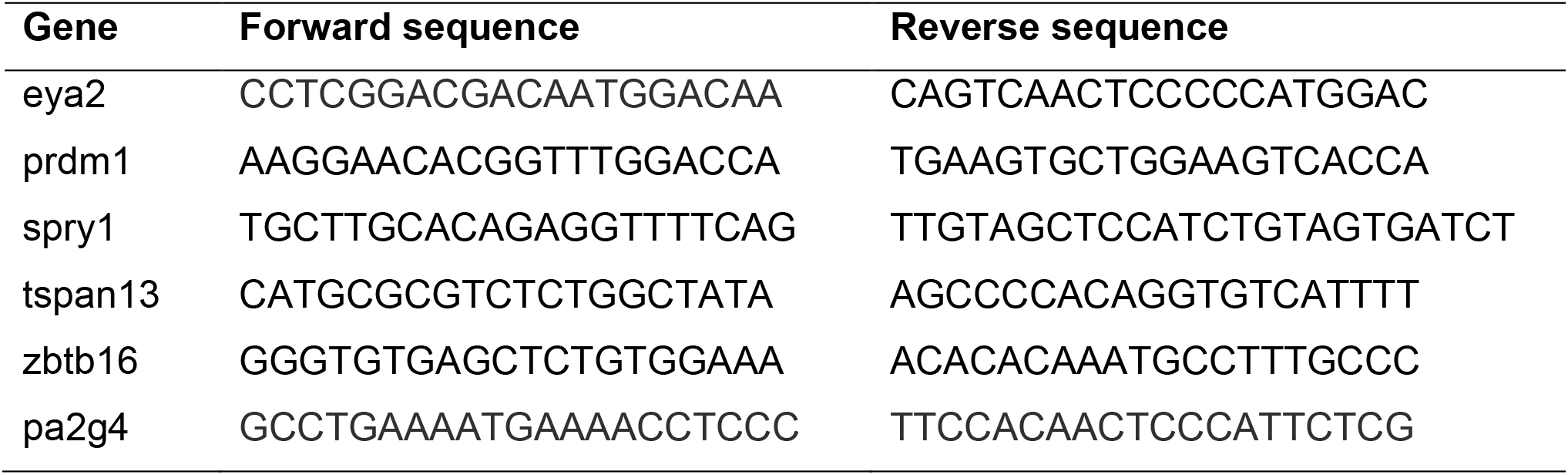
qPCR primer sequences.

## RESULTS

### Expression of several otic genes requires Six1

Several microarray, RNAseq and ChIPseq assays have identified a large number of potential targets of Six1 transcriptional regulation [38–42]. We chose five putative targets from these lists based on their expression in the otic vesicle, and one novel cofactor that interferes with Six1-Eya1 transcriptional activation [43]. First, we tested whether the expression of these genes in the otic vesicle (OV) is altered when endogenous Six1 protein is knocked-down by injecting a previously validated mixture of two translation-blocking antisense morpholino oligonucleotides (Six1MOs) [21] into the neural crest/placode precursor blastomeres on one side of the embryo. The expression domain and/or intensity of each gene was reduced in the majority of embryos (*eya2*, 83.1%, n=124; *prdm1*, 96.7%, n=60; *spry1*, 62.2%, n=82; *tspan13*, 85.1%, n=47; *zbtb16*, 72.1%, n=61; *pa2g4*, 78.4%, n=51; Fig. 1). These results confirm that each gene requires an adequate level of Six1 protein for their normal expression in the OV.

**Figure 1:**
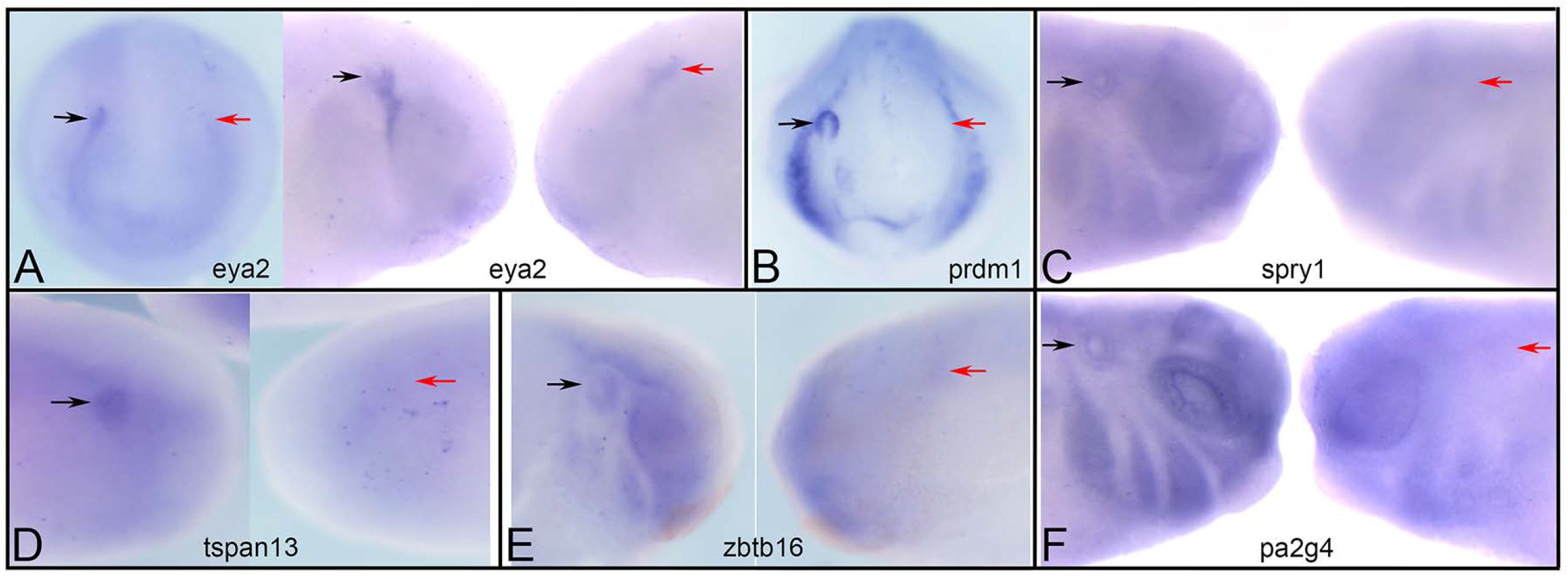
Six1 is required for otic gene expression. The otic placode (**A**, left image) or otic vesicle (**A**, right image, **B-F**) expression of: *eya2* (**A**), *prdm1* (**B**), *spry1* (**C**), *tspan13* (**D**), *zbtb16* **(E)** and *pa2g4* (**F**) was reduced on the MO-mediated knock-down side of each embryo (red arrow) compared to the control side (black arrow) of the same embryo. A, left image and B are frontal views; A, right image and C-F are side views. Dorsal is to the top.

Since Six1 can act as either a transcriptional activator or repressor depending upon the presence of different co-factors [21, 65–67], we tested whether the putative target genes were activated (either directly or indirectly) by Six1 by injecting *Six1VP16* mRNA, or repressed (either directly or indirectly) by Six1 by injecting *Six1EnR* mRNA, and comparing the frequency of the phenotypes to that resulting from expressing SixWT. For *eya2*, Six1WT-150 (150pg of mRNA), Six1WT-400 (400pg of mRNA) and Six1EnR (100pg of mRNA) reduced its expression at the same frequencies (p>0.05), whereas Six1VP16 (100pg of mRNA) caused a significantly lower frequency of repression and expanded expression in about 25% of the embryos (Fig. 2A-C). Since the Six1VP16 phenotype frequencies were significantly different from Six1WT (p<0.0001), whereas the Six1EnR frequencies were not, we conclude that Six1WT acts to repress *eya2* expression in the embryo. Similar results were observed for *prdm1* (p<0.00001; Fig. 2D-F). For *spry1*, Six1VP16 and Six1EnR both repressed its expression at frequencies indistinguishable from either dose of Six1WT (Fig. 2G-I), indicating multiple pathways for its regulation by Six1. For *tspan13*, Six1VP16 was significantly more repressive compared to Six1WT-150pg (p<0.05) but not different compared to Six1WT-400pg; Six1EnR repressed *tspan13* at frequencies indistinguishable from either Six1WT dose (Fig. 2J-L). These results suggest that the endogenous levels of Six1 in the embryo differentially regulate *tspan13* expression: at high levels, it reduced *tspan13* expression via transcriptional activation, whereas at low levels it reduced *tspan13* expression via transcriptional repression. For *pa2g4*, Six1VP16 repressed its expression at a frequency indistinguishable from Six1WT-150, whereas Six1EnR repressed at a similar frequency as Six1WT-400 (Fig. 2M-O). These results suggest that the endogenous levels of Six1 in the embryo differentially regulate *pa2g4* expression, but in contrast to *tspans13*, at high levels, Six1 reduced *pa2g4* by transcriptional repression and at low levels it reduced *pa2g4* via transcriptional activation.

**Figure 2:**
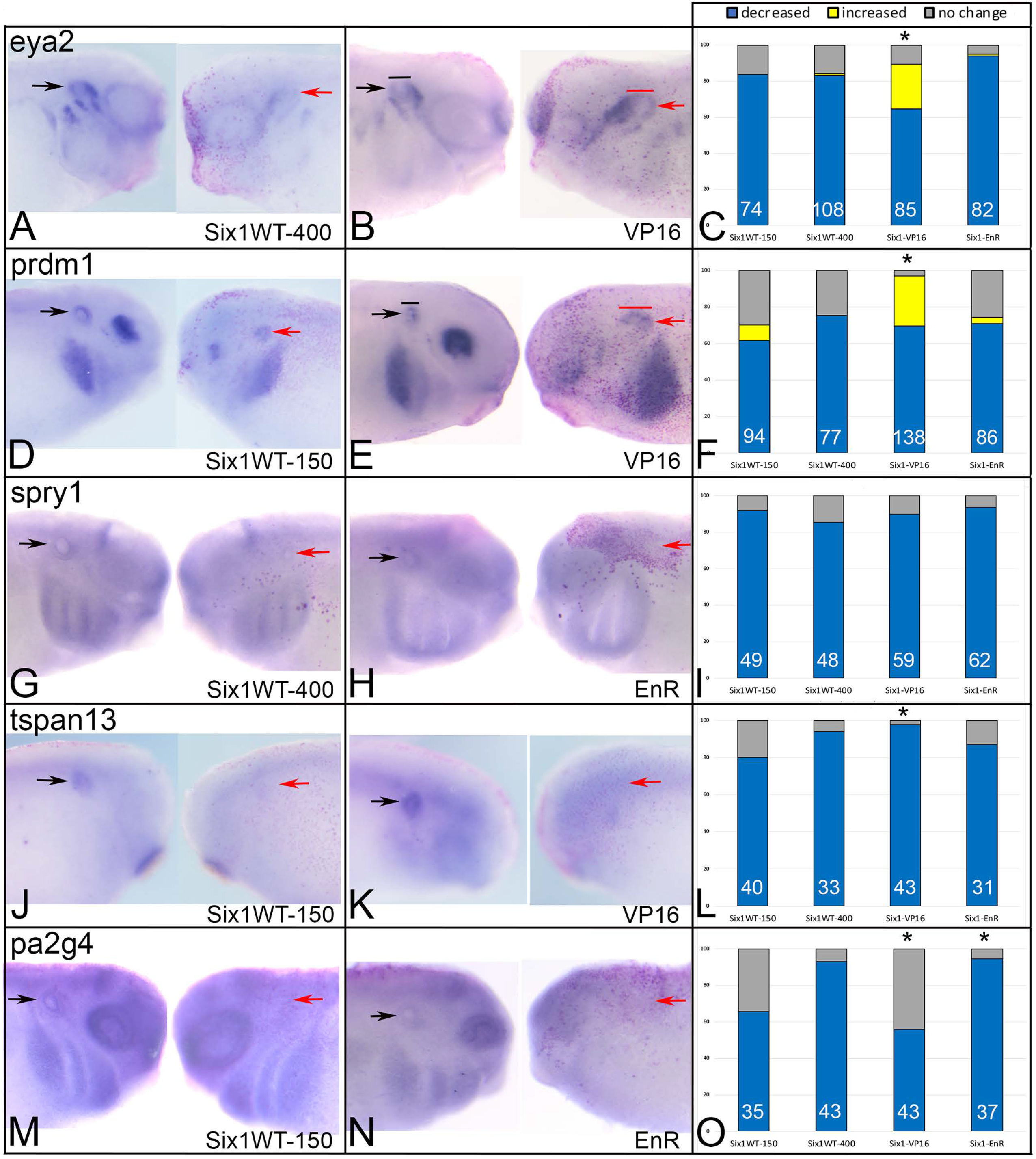
Increased Six1 alters otic gene expression. Transcripts encoding wild-type Six1 (Six1WT, either 150 or 400 pg), activating Six1 (VP16, 100pg) or repressive Six1 (EnR, 100pg) were microinjected into neural crest and cranial placode precursor blastomeres on one side of the embryo. Gene expression was assayed by ISH for *eya2* (**A, B**), *prdm1* (**D, E**), *spry1* (**G, H**), *tspan13* (**J, K**) and *pa2g4* (**M, N**). The control, uninjected side of each embryo is on the left and the mRNA-injected side of the same embryo is on the right. Otic gene expression on the control side is indicated by black arrows, and that on the mRNA-injected side by red arrows. In B and E, the width of the otic vesicle is indicated by a black (control) or red (injected) bar. Frequencies of the effects (blue, decreased expression; yellow, increased expression; grey, no change) are indicated by bar graphs (**C, F, I, L, O**). For *eya2* (**C**) and *prdm1* (**F**), the frequencies for Six1VP16 were significantly different from both Six1WT-150 and SixWT-400 (*, *p*<0.0001). For *tspn13* (**L**), the frequencies for Six1VP16 were significantly different from Six1WT-150 (*, *p*<0.05), but not from SixWT-400. For *pa2g42* (**O**), the frequencies for Six1VP16 were significantly different from SixWT-400 (*, *p*<0.0001), and the frequencies for Six1EnR were significantly different from SixWT-150 (*, *p*<0.01). White numbers inside each bar denotes the number of embryos analyzed.

### BOR mutations differentially reduce cranial expression of putative Six1 targets

To determine whether some of the *SIX1* mutations found in BOR patients affect the expression of the investigated otic genes, we performed qPCR analysis of mRNAs extracted from whole larval heads (Fig. 3). For *eya2*, both Six1WT-150 and V17E significantly reduced expression; however, there was no significant difference between them indicating V17E was as active as Six1WT. At 400 pg, Six1WT, W122R and Y129C were not significantly different from control or each other, indicating that W122R and Y129C were as active as Six1WT. Interestingly, R110W reduced *eya2* expression significantly more than Six1WT, suggesting that it may be dominant-repressive. For *prdm1*, V17E significantly reduced expression compared to control and Six1WT-150. At 400pg, R110W and W122R significantly reduced *prdm1* expression below control levels, whereas Six1WT-400 and Y129C were not significantly different from control. For *spry1*, V17E significantly reduced expression below control levels, but was indistinguishable from Six1WT-150. At 400pg, there were no significant differences between controls, Six1-WT and mutants. For *tspan1*, there were no significant differences between controls, Six1WT-150 and V17E; however, at 400pg R110W significantly reduced *tspan1* expression compared to control and those of Six1WT, W122R and Y129C. For *zbtb16*, there were no significant differences between controls, either level of Six1WT or any of the mutants. For *pa2g4*, Six1WT-150 significantly reduced expression and V17E reduced expression significantly more than Six1WT-150. At 400pg, Six1WT, R110W and W122R significantly reduced *pa2g4* levels compared to control. Moreover, R110W reduced levels significantly more than Six1WT and Y129C.

**Figure 3:**
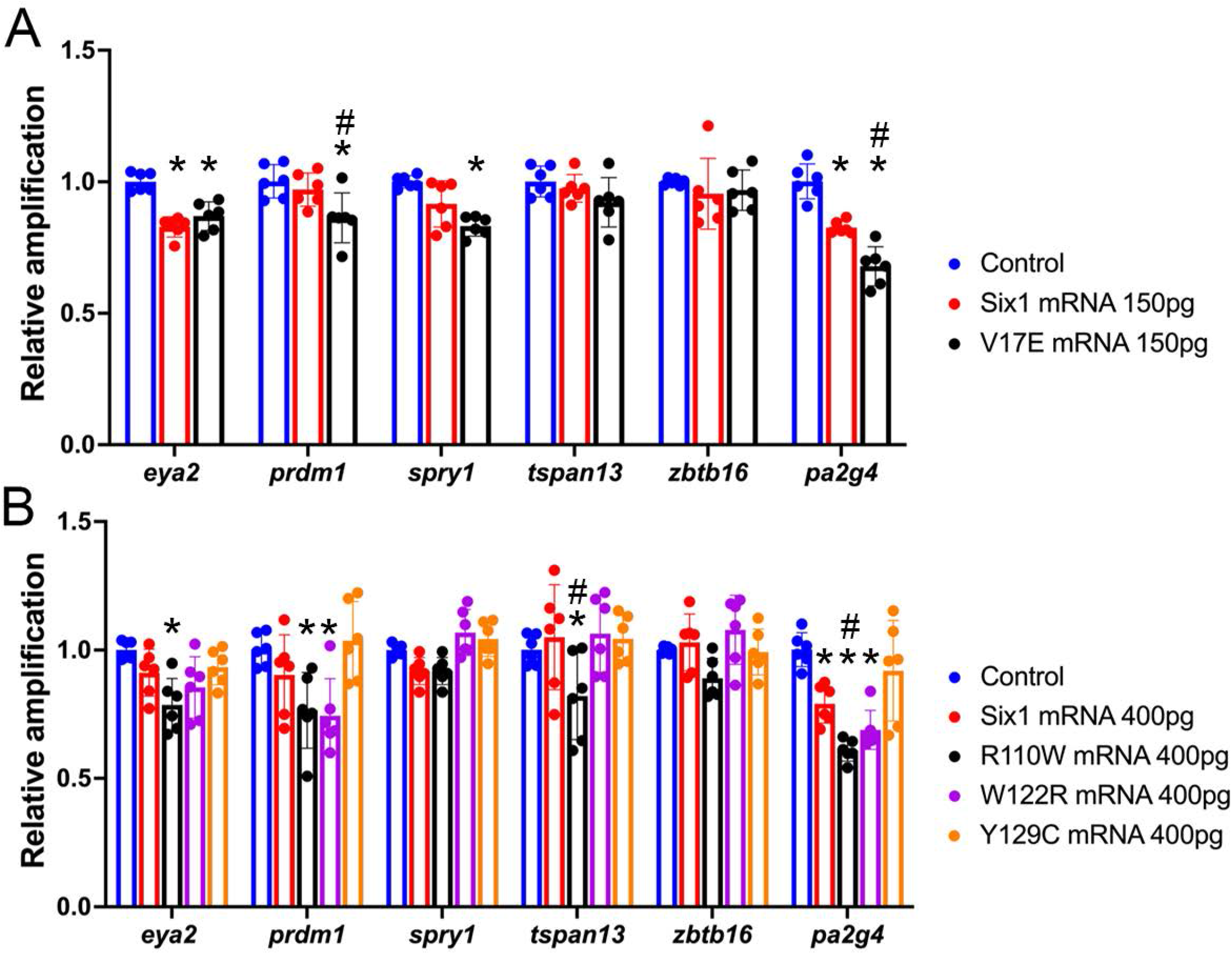
Gene expression changes in whole head samples assessed by qPCR. **A.** Levels of expression in whole heads collected from uninjected, sibling-matched embryos (controls, blue bars), and embryos injected with mRNA encoding either Six1WT (red bars) or V17E (black bars). Significant differences from control (*p*<0.05) are indicated by an asterisk; significant differences from Six1WT (*p*<0.05) are indicated by #. **B.** Levels of expression in whole heads collected from uninjected, sibling-matched embryos (controls, blue bars), and embryos injected with mRNA encoding either Six1WT (red bars), R110W (black bars), W122R (purple bars) or Y129C (orange bars). Significant differences from control (*p*<0.05) are indicated by an asterisk; significant differences from Six1WT (*p*<0.05) are indicated by #. Data are from three independent samples run in duplicate.

Overall, these results indicate that increased expression of Six1WT significantly reduced the expression in whole heads of some (*eya2, pa2g4*) but not all of the measured genes, and that only some of the mutants have effects comparable to Six1WT. V17E significantly reduced the expression in whole heads of most genes (*eya2, prdm1, spry1, pa2g4*), sometimes significantly more than a comparable dose of Six1WT (*prdm1, pa2g4*). R110W significantly reduced the expression in whole heads of *eya2, prdm1, tspan13* and *pa2g4*, but this was only significantly different from Six1WT-400 for *tspan13* and *pa2g4*. W122R significantly reduced the expression in whole heads of *prdm1* and *pa2g4*, but they were not significantly different from Six1WT-400. Y129C did not significantly reduce any of the measured genes compared to controls, indicating that is does not have the same activity as Six1WT. Thus, each SIX1 mutant showed differential effects on the cranial expression levels of these genes.

### 3.3 BOR mutations differentially affect otic gene expression

Our previous work demonstrated that these four BOR mutants have differential effects on the expression of neural border, neural crest and PPE genes, which ultimately results in changes in the expression of a few genes that pattern the OV (*dlx5, irx1, otx2, pax2 sox9*) [14]. Here, we focused on Six1 putative targets and co-factors.

At 150pg, V17E reduced the otic expression of *eya2, tspan13, zbtb16* and *pa2g4* at a frequency indistinguishable from Six1WT-150 (Fig. 4A, G, I, K), whereas it reduced *prdm1* more frequently and *spry1* less frequency compared to Six1WT-150 (Fig. 4C, E). At 150pg, R110W and W122R reduced the otic expression of all 6 genes at a significantly lower frequency compared to Six1WT-150. At 150pg, Y129C reduced the otic expression of *eya2, prdm1, tspan13*, and *zbtb16* at a significantly lower frequency than Six1WT-150, but showed no difference for *spry1* or *pa2g4*. We also compared the frequencies between the four mutants at 150pg (Fig. 4A, C, E, G, I, K). For *spry1*, the four mutants caused repression at a similar frequency. For *eya2, prdm1, tspan13* and *zbtb16*, V17E caused significantly more repression of gene expression compared to R110W, W122R and Y129C. For *eya2* and *prdm1*, there were no significant differences between R110W, W122R and Y129C. For *tspan13*, W122R was significantly different from R110W and Y129C was significantly different from W122R. For *zbtb16*, W122R and Y129C were significantly different from R110W. For *pa2g4*, V17E caused significantly more repression compared to R110W and W122R; the effects of V17E and Y129C were indistinguishable. Occasionally the otic expression of *eya2, prdm1* and *tspan13* were broader due to R110W, W122R or Y129C, but this was not observed for V17E.

**Figure 4:**
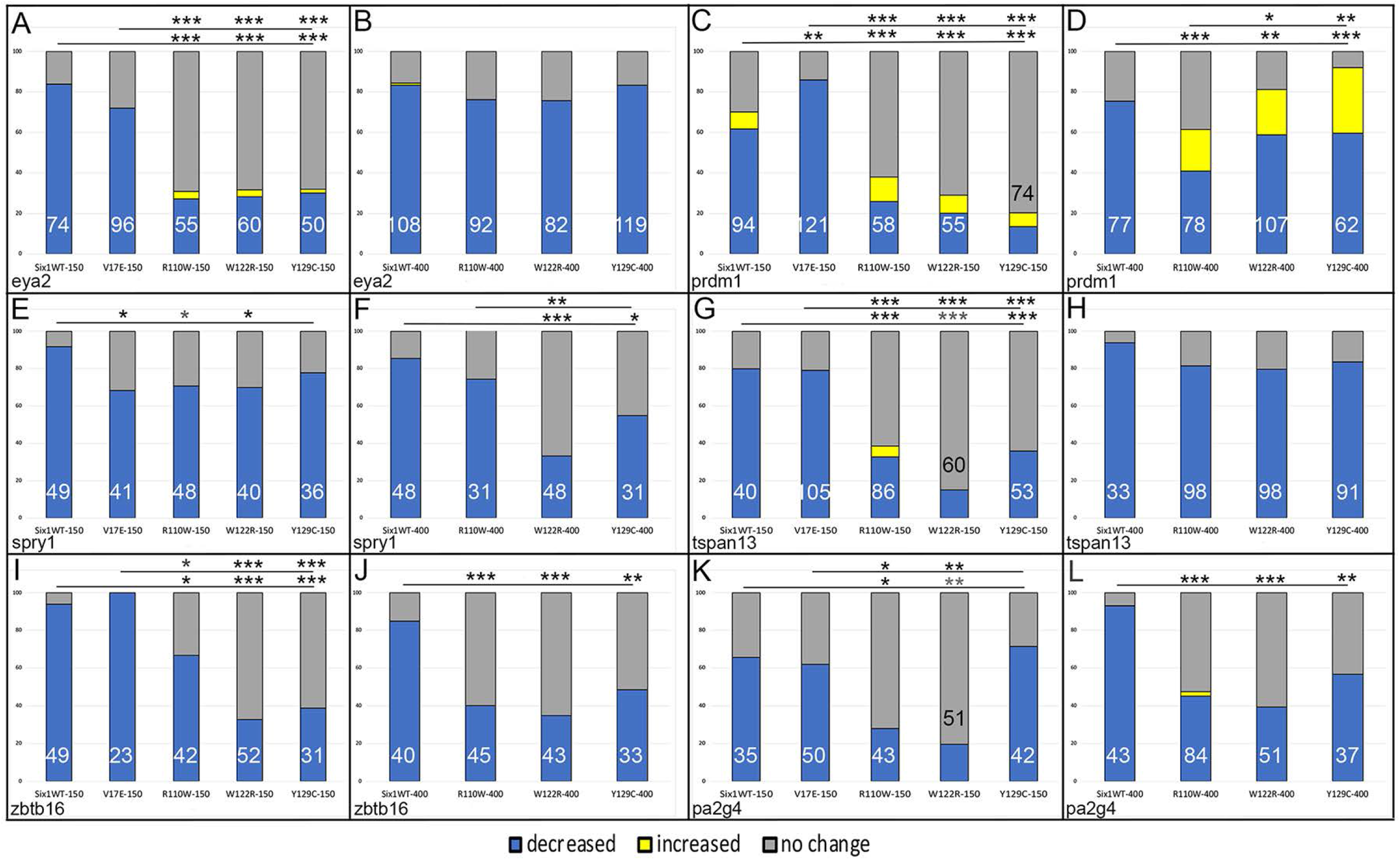
Otic gene expression is altered by Six1 mutants as assessed by ISH. **A:** Percentages of embryos in which *eya2* otic expression was decreased (blue), increased (yellow) or not changed (grey) after microinjection of 150pg of Six1WT, V17E, R110W, W122R or Y129C mRNAs. Numbers inside bars denote number of embryos analyzed. Lower line and asterisks indicate comparison of Six1WT frequencies to those of each mutant. Upper line and asterisks indicate comparison of V17E frequencies to those of the other three mutants. *, *p*<0.01; **, *p*<0.001; ***, *p*<0.0001 comparison by chisquare test. **B.** Percentages of embryos in which *eya2* otic expression was altered after microinjection of 400pg of Six1WT, R110W, W122R or Y129C mRNAs. There were no significant differences between the different groups. **C.** Percentages of embryos in which *prdm1* otic expression was altered after microinjection of 150pg of Six1WT, V17E, R110W, W122R or Y129C mRNAs. Labeling as in A. **D.** Percentages of embryos in which *prdm1* otic expression was altered after microinjection of 400pg of Six1WT, R110W, W122R or Y129C mRNAs. Labeling as in A. **E**. Percentages of embryos in which *spry1* otic expression was altered after microinjection of 150pg of Six1WT, V17E, R110W, W122R or Y129C mRNAs. Labeling as in A. **F.** Percentages of embryos in which *spry1* otic expression was altered after microinjection of 400pg of Six1WT, R110W, W122R or Y129C mRNAs. Labeling as in A. **G.** Percentages of embryos in which *tspan13* otic expression was altered after microinjection of 150pg of Six1WT, V17E, R110W, W122R or Y129C mRNAs. Labeling as in A. **H.** Percentages of embryos in which *tspan13* otic expression was altered after microinjection of 400pg of Six1WT, R110W, W122R or Y129C mRNAs. There were no significant differences between the different groups. **I.** Percentages of embryos in which *zbtb16* otic expression was altered after microinjection of 150pg of Six1WT, V17E, R110W, W122R or Y129C mRNAs. Labeling as in A. **J.** Percentages of embryos in which *zbtb16* otic expression was altered after microinjection of 400pg of Six1WT, R110W, W122R or Y129C mRNAs. Labeling as in A. **K.** Percentages of embryos in which *pa2g4* otic expression was altered after microinjection of 150pg of Six1WT, V17E, R110W, W122R or Y129C mRNAs. Labeling as in A. **L.** Percentages of embryos in which *pa2g4* otic expression was altered after microinjection of 400pg of Six1WT, R110W, W122R or Y129C mRNAs. Labeling as in A.

As previously reported [14], embryos do not survive a microinjection of >150pg of V17E, whereas R110W, W122R and Y129C are tolerated at a higher concentration. At 400pg, R110W reduced the otic expression of *eya2, spry1* and *tspan13* at the same frequency as Six1WT-400 (Fig. 4B, F, H), whereas it reduced *prdm1, zbtb16* and *pa2g4* otic expression significantly less frequently than Six1WT-400 (Fig. 4D, J, L). W122R and Y129C also reduced the otic expression of *eya2* and *tspan13* at the same frequency as Six1WT-400, whereas they reduced *prdm1, spry1, zbtb16* and *pa2g4* otic expression significantly less frequently than Six1WT-400. We also compared the frequencies between the three mutants at 400pg (Fig. 4B, D, F, H, J, L). For *eya2, tspan13, zbtb16* and *pa2g4*, all 3 mutants reduced gene expression at similar frequencies. For *prdm1*, R110W was significantly less repressive than W122R or Y129C, whereas W122R and Y129C frequencies were indistinguishable. For *spry1*, W122R was significantly less repressive than R110W or Y129C; R110W and Y129C frequencies were indistinguishable. Altogether, these results indicate that the four BOR mutants differentially affect otic gene expression. For many of the monitored genes, the mutants were significantly less repressive than Six1WT, and in a few cases some mutants, particularly *prdm1*, expanded otic gene expansion.

## DISCUSSION

The Six1 homeodomain containing transcription factor plays a key role in vertebrate cranial placode development [15–19]. Loss-of-function by knock-down of endogenous protein or by genetic deletion results in reduced expression of placode genes and a variety of defects during otic development spanning otic placode formation to sensory hair cell differentiation [20–33]. Mutations in human *SIX1* underlie some cases of BOR, but because there is considerable variability in the presence, type and severity of craniofacial and renal abnormalities even among family members carrying the same mutation, it has not been possible to associate a specific mutation with a particular suite of clinical presentations [9–12,36,37]. One way to begin to address how each mutation might result in a particular phenotype is to study how the variant protein affects developmental processes in an experimental animal model. To this end, we introduced four of the human mutations into *Xenopus Six1* and found that expressing each mutant protein in a wild-type embryo had a different suite of effects on neural border, neural crest, PPE and a few OV genes that subsequently resulted in subtle abnormalities in otic morphogenesis [14]. Herein, we expanded our analysis to OV genes that previously were reported by large-scale screens to be likely targets of Six1 and/or co-factors [38–43].

First, we confirmed that Six1 is required for otic expression of the selected targets/co-factors (Table 2). Next, to determine the mode by which Six1 might regulate their expression we expressed fusion constructs that render Six1 as either a transcriptional activator (Six1VP16) or a transcriptional repressor (Six1EnR). We found that expression of *eya2* and *prdm1* are reduced by Six1 via transcriptional repression, whereas expression of *spry1, tspan13*, and *pa2g4* are reduced by both transcriptional activation and repression, and in the cases of *tspan13* and *pa2g4*, the mode depends upon the level of Six1 (Table 2). These results indicate that Six1 likely regulates these genes by both direct and indirect mechanisms. Interestingly, ChIP analysis of the E10.5 mouse OV, which is similar to the OV stage studied herein, described Six1 binding sites in close proximity to *eya2, prdm1* and *spry1* genes [42], suggesting that at least some of these effects are likely mediated by direct binding to the target genes. Whether Six1 binding to these sites results in activation versus repression will need to be resolved in future studies.

**Table 2:** Summary of effects Six1 on otic vesicle gene expression.

|  | Six1 required | Does Six1 activate or repress? | V17E | R110W | W122R | Y129C |
| --- | --- | --- | --- | --- | --- | --- |
| *eya2* | yes | repress | head: ↓ = Six1WT<br>otic: ↓ = Six1WT | head: ↓ = Six1WT<br>otic: low: ↓ < Six1WT<br>high: ↓ = Six1WT | head: no reduction<br>otic: low: ↓ < Six1WT<br>high: ↓ = Six1WT | head: no reduction<br>otic: low: ↓ < Six1WT<br>high: ↓ = Six1WT |
| *prdm1* | yes | repress | head: ↓ > Six1WT<br>otic: ↓ > Six1WT | head: ↓ = Six1WT<br>otic: low: ↓ < Six1WT<br>high: ↓ < Six1WT | head: ↓ = Six1WT<br>otic: low: ↓ < Six1WT<br>high: ↓ < Six1WT | head: no reduction<br>otic: low: ↓ < Six1WT<br>high: ↓ < Six1WT |
| *spry1* | yes | both | head: ↓ = Six1WT<br>otic: ↓ < Six1WT | head: no reduction<br>otic: low: ↓ < Six1WT<br>high: ↓ = Six1WT | head: no reduction<br>otic: low: ↓ < Six1WT<br>high: ↓ < Six1WT | head: no reduction<br>otic: low: ↓ = Six1WT<br>high: ↓ < Six1WT |
| *tspan13* | yes | low: repress<br>high: activate | head: no reduction<br>otic: ↓ = Six1WT | head: ↓ > Six1WT<br>otic: low: ↓ < Six1WT<br>high: ↓ = Six1WT | head: no reduction<br>otic: low: ↓ < Six1WT<br>high: ↓ = Six1WT | head: no reduction<br>otic: low: ↓ < Six1WT<br>high: ↓ = Six1WT |
| *zbtb16* | --- | --- | head: no reduction<br>otic: ↓ = Six1WT | head: no reduction<br>otic: low: ↓ < Six1WT<br>high: ↓ < Six1WT | head: no reduction<br>otic: low: ↓ < Six1WT<br>high: ↓ < Six1WT | head: no reduction<br>otic: low: ↓ < Six1WT<br>high: ↓ < Six1WT |
| *pa2g4* | yes | low: activate<br>high: repress | head: ↓ > Six1WT<br>otic: ↓ = Six1WT | head: ↓ > Six1WT<br>otic: low: ↓ < Six1WT<br>high: ↓ < Six1WT | head: ↓ = Six1WT<br>otic: low: ↓ < Six1WT<br>high: ↓ < Six1WT | head: no reduction<br>otic: low: ↓ = Six1WT<br>high: ↓ < Six1WT |

**Legend:** ↓ = Six1WT, reduced the same as Six1WT; ↓ > Six1WT, reduced more compared to Six1WT; ↓ < Six1WT, reduced less compared to Six1WT; ---, data reported in a pending, unpublished manuscript.

Several studies have investigated the biochemical effects of various BOR mutations. For example, a study of *EYA1* mutations concluded that the variant proteins function in a dominant-negative fashion [68]. Based on cell culture experiments, SIX1 mutant proteins also have been suggested to act as dominant-negative by competing with wild-type protein for DNA binding sites or by competing for the EYA1 co-factor [2,69]. With this information in hand, we sought to understand the gene expression phenotypes caused by four of the SIX1 mutants when they are expressed in the embryo. To address whether BOR mutant versions of Six1 have the same activity in the embryo as Six1WT, we first assayed gene expression in dissected heads by qPCR. We observed that V17E, R110W, and W122R, like Six1WT, significantly reduced the expression of many of the selected genes, sometimes significantly more than a comparable dose of Six1WT (Table 2). Interestingly, *zbtb16* expression was not altered by any of the mutants and Y129C did not alter the expression of any of the selected genes. Although these data demonstrate that each SIX1 mutant had a different combination of effects on the cranial expression levels of the selected genes, the assay lacks tissue specificity: whole heads were collected. Therefore, we next expressed the mutants only in the NC and PPE precursors that contribute to the middle and inner ear and assessed OV gene expression by ISH.

The V17E mutation lies in the first α-helix of the SD. It interferes with the interaction between Six1 and Eya1/Eya2, reduces translocation of cytosolic Eya protein to the nucleus, disrupts binding to DNA due to steric hindrance and is transcriptionally deficient [13,14,70]. Our previous work showed that V17E caused the same effects as low levels of Six1WT on neural border gene expression, as well as some NC, PPE and OV genes. However, it reduced the expression of *sox9* in the neural crest and otic placode less than Six1WT. In contrast, it reduced *irx1* in the PPE more than Six1WT, reduced *sox11* in the PPE whereas Six1WT expanded *sox11*, and had differential effects on OV genes. *pax2* was reduced less by V17E than by Six1WT, whereas *sox9, tbx1* and *dlx5* were reduced more than by Six1WT, and V17E reduction of *irx1* and *otx2* expression were similar to Six1WT [14]. Of these otic genes, there is only evidence for *irx1* to be a direct target of Six1 [33,42], so we investigated the effects on additional otic genes that are likely to be targets and/or co-factors. Similar to Shah et al [14], V17E reduced one otic gene less (*spry1*) and one otic gene (*prdm1*) more compared to Six1WT, but most effects were indistinguishable from Six1WT (Table 2). Since increased Six1 protein predominantly acts as a transcriptional repressor of these genes (Fig. 2; Table 2), V17E likely has a hypomorphic effect on *spry1* and a dominant-negative effect on *prdm1*, whereas it is as effective as Six1WT for the other genes.

The R110W mutation lies in the sixth α-helix of the SD. It reduces the interaction with EYA1 but not with EYA2 [10,13]. It can transport EYA2 and Eya1 to the nucleus [13,14] and bind to DNA [10,13], but does not activate transcription in a luciferase reporter assay in the presence of Eya1 or EYA2 [10,13,14]. Our previous work showed that R110W reduced neural border, NC and PPE gene expression significantly less than Six1WT and sometimes caused them to be broader. R110W (400pg) reduced the otic expression of *pax2, sox9, tbx1*, and *dlx5* significantly less than Six1WT (400pg), whereas the reduction of *irx1* and *otx2* expression was similar to that of Six1WT [14]. In the present study, we assessed the effects of both a low dose (150pg) and high dose (400pg) of Six1WT and mutants. At 150pg, R110W reduced each otic gene significantly less frequently than Six1WT; at 400pg, it reduced *prdm1, zbtb16* and *pa2g4* significantly less frequently. Together, these studies indicate that the R110W variation renders Six1 less effective at repressing target genes. Since R110W can bind Eya1 and transport it to the nucleus, we posit that this deficiency may be due to interference with the binding of co-factors that mediate transcriptional repression, such as Groucho/TLE, Pa2G4 and Mcrs1 [43,56,71].

The W122R mutation, located in the linker region between the sixth α-helix and the HD, has been postulated to affect DNA binding efficiency [13]. Although it can translocate Eya1 to the nucleus in HEK293T cells, it cannot activate transcription in the presence of Eya1 [14]. Our previous work showed that W122R reduced neural border, NC and PPE gene expression significantly less than Six1WT and sometimes caused them to be broader. W122R (400pg) reduced the otic expression of *pax2* and *sox9* significantly less than Six1WT (400pg), whereas the reduction of *tbx1, dlx5, irx1* and *otx2* expression were similar to that of Six1WT [14]. In the present study, at 150pg W122R reduced all 6 otic genes significantly less frequently than Six1WT, and at 400pg, it reduced *prdm1, spry1, zbtb16* and *pa2g4* significantly less frequently. Since these results are very similar to those of R110W, we predict that its effects are due to deficits in binding repressive co-factors rather than deficits in DNA binding.

The Y129C mutation, located in the N-terminal region of the HD, can interact with Eya1/EYA2 and translocate them to the nucleus, but it does not bind to DNA or activate transcription in a luciferase reporter assay in the presence of Eya1 or EYA2 [10,13,14]. Our previous work showed that Y129C caused the same effect as Six1WT on neural border gene expression [14], but it reduced NC and PPE gene expression significantly less than Six1WT and sometimes caused them to be broader. Y129C (400pg) reduced the otic expression of *pax2* and *sox9* significantly less than Six1WT (400pg), whereas the reduction of *tbx1, dlx5, irx1* and *otx2* expression was similar to that of Six1WT [14]. In the present study, at 150pg Y129C reduced the otic expression of *eya2, prdm1, tspan13* and *zbtb16* significantly less frequently than Six1WT, and at 400pg, it reduced *prdm1, spry1, zbtb16* and *pa2g4* significantly less frequently. If Y129C acted as a dominant-negative by reducing the available endogenous Eya1, as has been postulated [2,69], we predict it would cause endogenous Six1WT to bind to target genes without Eya1 and thus be more repressive. Accordingly, perhaps Y129C, as proposed for W122R, binds to and thereby reduces the endogenous availability of repressive co-factors.

### Conclusion

The expression of each of the four mutant Six1 proteins in wild-type embryos has identified different effects on developmental precursor populations that give rise to the craniofacial tissues that are dysmorphic in BOR patients. By focusing on otic genes that are likely to be direct targets and/or co-factors of Six1, we demonstrate that each mutant tends to show a lower frequency of gene expression reduction compared to Six1WT. Since Six1 acts as a transcriptional repressor in the absence of Eya1, and given that R110W, W122R and Y129C can bind Eya1 and transport it to the nucleus, one likely cause of these mutants showing a less reduced phenotype is that they do not bind to repressive co-factors. Alternatively, altered *pa2g4* expression may account for these changes. Since Pa2G4 is a repressive co-factor that competes with Eya1 binding [43], the lower frequency of reduced *pa2g4* would result in higher levels of its protein being available. It is interesting that V17E tends to have effects that are significantly different from R110W, W122R and Y129C, suggesting that the underlying cause is its inability to bind and translocate Eya proteins. An important next step in understanding these changes in gene expression is to identify the function of each of the putative candidate cofactors [55] and to identify where they bind and whether they interfere with Eya protein binding.

## Funding

NIH-NIDCR R01 DE022065

NIH-NIDCR R01 DE026434

## ACKNOWLEDGEMENTS

This work was made possible with the support of Xenbase (http://www.xenbase.org/, RRID: SCR_003280) and the National *Xenopus* Resource (http://mbl.edu/xenopus/, RRID:SCR_013731). We thank Professor Charles Sullivan and Ms. Jenni Xu for performing some of the microinjections. This research was funded by the National Institutes of Health (DE022065 and DE026434 to S.A.M) and the George Washington University Center for Undergraduate Fellowships and Research (to T.M. and S.A.).

